# Enamel Palaeoproteomics Successfully Distinguishes between Denisovans and European Neandertals

**DOI:** 10.64898/2026.08.22.746388

**Authors:** Paula Kotli

## Abstract

Ancient DNA (aDNA) has transformed the study of hominin relationships, but its preservation in ancient fossils is often limited. Enamel palaeoproteomics offers an alternative molecular approach for taxonomic analysis. In this study, we re-analyse published DDA mass spectrometry data ^1^ from the Denisovan-attributed Penghu 1 mandible (PXD054412) ^2^ and a Neandertal enamel specimen from Gruta de Oliveira, Portugal (PXD038154) ^3^. Five AMBN peptides carrying the Valine-273 substitution (V273) were validated in the Penghu 1 enamel, three of which were independently detected in both DDA acquisitions. No V273-containing peptide signal was detected in the Neandertal dataset. Conversely, the ancestral Methionine-273 peptide REDPM[+16]AYG was detected exclusively in the Neandertal specimen. Extracted ion chromatograms, isotopic envelope confirmation, and MS2 fragmentation spectra, all support the reported peptide assignments. AMELY-specific peptides additionally support male sex assignment for both ancient individuals. Together, these results confirm the taxon-specific mutual exclusivity of AMBN V273 and M273 variants across Denisovan and Neandertal lineages, establishing AMBN M273V as a molecularly validated diagnostic marker for Denisovan identification from dental enamel. More broadly, targeted MS1 reanalysis of public proteomics datasets provides a scalable complement to ancient genomics for resolving hominin lineage identity and sex determination when DNA is not preserved.

## Introduction

Ancient dental enamel preserves endogenous proteins over timescales spanning hundreds of thousands to over one million years, making it one of the most valuable substrates for deep-time palaeoproteomics ^4,5^. The analysis of ancient enamel proteomes has emerged as a powerful complement to ancient DNA (aDNA) methods for phylogenetic reconstruction of archaic hominins, particularly in depositional contexts, including tropical, subtropical, and cave environments at low latitudes, where elevated temperatures preclude aDNA survival ^2,6^.

A particularly powerful refinement of palaeoproteomics is the exploitation of single amino acid variants (SAAVs): taxon-specific amino acid substitutions at defined positions in enamel proteins that allow specimen assignment to particular hominin lineages without full genome sequencing. This approach was foundational to the identification of *Homo antecessor* as a basal hominin related to the last common ancestor of Neandertals, Denisovans, and modern humans ^5^, and to the attribution of the Xiahe mandible to Denisovans via Collagen I SAAVs ^7^.

Denisovans present a unique challenge for palaeoproteomics: confirmed Denisovan specimens are extremely rare, and genomic data characterising their biology were initially derived from a single distal phalanx at Denisova Cave, Siberia ^8^, with subsequent discoveries remaining geographically and numerically limited. Their morphological boundaries remain poorly defined and their full geographic range incompletely characterised. Several candidate Denisovan specimens have been proposed on anatomical grounds, including the Penghu 1 mandible from Taiwan ^2^ and the Cobra Cave molar from northern Laos ^9^, but taxonomic assignment requires independent molecular evidence. In the absence of recoverable aDNA, as is the case for many East and Southeast Asian Pleistocene fossils, enamel proteomics represents the most promising tool for taxonomic validation.

Zanolli et al. ^10^ first predicted that a Valine residue at position 273 of Ameloblastin (AMBN M273V; rs564905233) constitutes a derived variant specific to Denisovans, based on comparative analysis of tooth protein polymorphisms from Neandertal and Denisovan genome sequences against modern human and non-human primate data. This AMBN position exhibits a Methionine residue (M273) in modern humans, Neandertals, and all non-human primates examined, while the Valine variant (V273) is observed in Denisova 3 and at very low frequency in most modern human populations ^10,11^. Importantly, Zanolli et al. ^10^ also predicted a Neandertal -specific variant, AMBN G78S (rs143795139), in the Vindija 33.15 specimen, underscoring the potential of AMBN as a multi-variant diagnostic protein for archaic hominin identification, see aligments at Fig.3.

The recent characterisation of the Penghu 1 mandible by Tsutaya et al. ^2^ provided the first direct proteomic evidence for AMBN V273 in a Pleistocene fossil, identifying from the acid-etched enamel fraction at least 17 PSMs covering position 273. This finding supported the assignment of Penghu 1 as a probable Denisovan individual and confirmed the value of AMBN as a diagnostic target. However, PSM-level identification, while informative, does not provide the quantitative rigour of targeted proteomics approaches. Extracted ion chromatograms (XIC), isotopic envelope verification, and MS2 fragmentation spectra are required to confirm peptide identity at the highest confidence level and to rule out isobaric interference, spectral misassignment, or false positives arising from database search artefacts ^12^. Furthermore, demonstrating the specificity of the assay, the absence of V273 in a non-Denisovan hominin, requires comparative analysis across taxa within a single validated workflow ^6^.

Here, we present a re-analysis of DDA data from the Penghu 1 specimen (PXD054412) alongside data from a Neandertal enamel sample from Gruta de Oliveira, Portugal (PXD038154; Shaw et al. ^3^). Five V273 containing AMBN peptides were validated by us in the Denisovan specimen across two independent DDA data acquisitions, the ancestral M273 peptide exclusively detected in the Neandertal specimen, and male sex confirmed in both individuals via AMELY specific peptide detection. Together, these analyses provide the first targeted quantitative validation of AMBN M273V as a proteomic diagnostic marker for Denisovans and provide a reproducible assay framework applicable to future Pleistocene enamel specimens.

## Experimental Section

### Spectral data sources

DDA data raw data for the Penghu 1 specimen were obtained from the ProteomeXchange Consortium repository under accession **PXD054412** 2. Two data-dependent acquisition (DDA) runs were analysed from the acid-etched enamel fraction:

- **Denisovan**_184_: 20221026_EXPL5_nLC3_GT_collab_77min_DDA_Enamel_184
- **Denisovan**_205_: 20221115_EXPL5_nLC3_GT_collab_77min_DDA_Enamel_205

These represent two independent DDA data acquisitions from the same Penghu 1 specimen. The acid-etching extraction method generates peptides without enzymatic digestion, preserving specimen integrity while yielding enamel-derived peptides suitable for downstream targeted DDA data analysis ^2,12^. DDA data for the Neandertal specimen (mandibular P_3_, Gruta de Oliveira, Torres Novas, Portugal; ∼90.1–92.0 ka BP) were obtained from ProteomeXchange under accession **PXD038154** 3, run identifier: **Neandertal _5001_1_A**.

Modern human dental enamel from one female and one male individual was included as an analytical control for sex determination assay validation (published data ^13^). Sex of the modern controls was confirmed biologically prior to sampling. These samples were processed using identical DDA data and Skyline parameters as the ancient specimens.

### Protein sequence database construction

A custom protein sequence database was constructed for initial Byonic database searches. The database incorporated AMBN sequence variants described by Zanolli et al. ^10^: the canonical human AMBN sequence (M273) and the Denisovan-specific variant (V273, corresponding to amino acid substitution M273V; rs564905233). Both sequence entries were included to allow simultaneous identification of V273- and M273-containing peptides within a single database search. AMBN sequence variants were adapted from Zanolli et al. ^10^ and entered as separate database entries. The database additionally included enamel and mineralized-tissue proteins observed in the analysed datasets, including AMELX, AMELY, ENAM, AMTN, DMP1, MGP, and CEMP1. AMELX and AMELY were included for enamel preservation assessment and sex determination, whereas AMBN M273/V273 peptides were the primary targets for taxonomic validation.

Database searches were performed in PMI-Byonic v5.11.4 (Protein Metrics Inc., Belmont, CA) using non-specific cleavage to accommodate the non-enzymatic acid-etching extraction (precursor mass tolerance: 5.0 ppm; fragment mass tolerance: 20.0 ppm; maximum missed cleavages: 3; protein FDR ≤1%). Variable modifications included deamidation of asparagine and glutamine (+0.9840 Da), oxidation of lysine, methionine, proline, and tryptophan (+15.9949 Da), phosphorylation of serine (+79.9663 Da), and dioxidation of methionine, proline, and tryptophan (+31.9898 Da). Byonic search results provided the initial peptide identification list and retention time used for Skyline targeted analysis. Only target PSMs, after exclusion of Reverse/decoy hits, were used for downstream protein-level summaries and deamidation calculations.

### Skyline targeted analysis

Targeted peptide quantification was performed in Skyline (64-bit, version 26.1.0.057) ^14^. A targeted peptide list was constructed comprising: (i) all predicted peptides spanning AMBN position 273 in both V273 (Denisovan) and M273 (ancestral) forms, including methionine-oxidised variants; (ii) AMELX-specific peptides for enamel protein preservation assessment; and (iii) AMELY-specific peptides for sex determination ^13,15,16^.

For each target peptide, doubly charged precursor ions were selected based on predicted *m/z* values. Extracted ion chromatograms (XICs) were generated for each target, and chromatographic peak areas were integrated using Skyline’s automated peak-picking with manual curation. Isotopic envelope confirmation was applied to all precursor ions with sufficient signal intensity. MS2 fragment ion spectra (b/y ion series) were extracted from available DDA spectra and matched to theoretical fragmentation patterns for each peptide sequence.

Retention time confirmation was used as an additional validation criterion: peptide identities were considered confirmed when observed retention times were consistent with Byonic identification anchors from the same dataset, and when replicate runs from the same specimen showed concordant retention times. Normalised peak areas are reported as the primary quantification metric.

A peptide was considered **validated** when all three of the following criteria were met: (1) a distinct XIC peak was present at the expected retention time; (2) the isotopic envelope of the precursor ion matched the theoretical distribution; and (3) MS2 fragmentation spectra confirmed ≥3 matching b- or y-ions for the peptide sequence.

A peptide was classified as **not detected (ND)** when no XIC peak was observed above background noise within a ±0.5 min retention time window centred on the expected retention time as showed by Byonic identification.

Deamidation of N and Q was calculated by in-house R-scripts.

## Results

### Detection and validation of Denisovan-associated AMBN V273 peptides

Five AMBN peptides carrying the Denisovan-associated Valine-273 (V273) substitution were detected and validated by Skyline. In the Penghu 1 enamel data (Table 1). All five peptides were validated in Denisovan_184_; three of the five were independently confirmed in Denisovan_205_. None of the five V273-containing peptides produced a validated signal in the Neandertal dataset (Table 1). The peptide **REDP-VAYG** (*m/z* 453.72, charge 2+; RT 19.48 min) yielded normalised areas of 8.11 × 10^9^ (Denisovan_184_) and 1.74 × 10^9^ (Denisovan_205_). The overlapping peptides **GREDPVA** (*m/z* 372.19, charge 2+; RT 11.2 min) and **REDPVAY** (*m/z* 425.21, charge 2+; RT 19.44 min) showed concordant signals in both runs, with areas of 9.15 × 10^9^ / 3.49 × 10^9^ and 1.10 × 10^10^ / 2.99 × 10^9^ in Denisovan_184_ / Denisovan_205_, respectively. The longer peptide **REDPVAYGAM[+16]FPGF** (*m/z* 786.86, charge 2+; RT 50.1 min; normalised area 3.54 × 10^8^) and the flanking peptide **EEVAGGREDPVA** (*m/z* 614.79, charge 2+; RT 18.65 min; normalised area 8.26 × 10^8^) were validated in Denisovan_184_ only, consistent with the higher overall signal intensity (Table 1 and Fig. 5). See also Fig. 3, which presents the alignment of AMBN sequences reconstructed manually in this analysis.

**Table 1.** AMBN peptides detected and validated by Skyline in Penghu 1 (Denisovan) and Gruta de Oliveira (Neandertal) enamel. Normalised peak areas reported. ND = not detected (no validated XIC peak within ± 0.5 min of expected retention time; see the Skyline targeted analysis subsection in Methods). Met-ox = methionine oxidation (+15.9949 Da). The Neandertal M273 area (bottom row) is shown in bold.

| Peptide sequence | Mod. | Precursor $m/z$ | RT (min) | AMBN variant | Denisovan<sub>184</sub><br>XIC | Denisovan<sub>205</sub><br>XIC | Neandertal<br>XIC |
| --- | --- | --- | --- | --- | --- | --- | --- |
| GREDPVA | — | 372.19 (2+) | 11.2 | V273 | $9.15 \times 10^9$ | $3.49 \times 10^9$ | ND |
| REDPVAY | — | 425.21 (2+) | 19.44 | V273 | $1.10 \times 10^{10}$ | $2.99 \times 10^9$ | ND |
| REDPVAYG | — | 453.72 (2+) | 19.48 | V273 | $8.11 \times 10^9$ | $1.74 \times 10^9$ | ND |
| REDPVAYGAM[+16]FPGF | Met-ox | 786.86 (2+) | 50.1 | V273 | $3.54 \times 10^8$ | ND | ND |
| EEVAGGREDPVA | — | 614.79 (2+) | 18.65 | V273 | $8.26 \times 10^8$ | ND | ND |
| REDPM[+16]AYG | Met-ox | 477.70 (2+) | 10.9 | M273 (ancestral) | ND | ND | **$2.22 \times 10^7$** |

Consistent retention times across both Penghu 1 acquisitions for the three co-detected peptides (GRED-PVA, REDPVAY, REDPVAYG) support reproducible peptide identification and reduce the likelihood of chromatographic artefacts. XIC profiles, isotopic envelope confirmation, and MS2 fragmentation evidence for the three highest-intensity V273 peptides are shown for Denisovan_184_ and Denisovan_205_ in Figures 1 and 2. The corresponding ancestral M273 peptide detected in the Neandertal dataset is shown in Figure 4.

**Figure 1.**
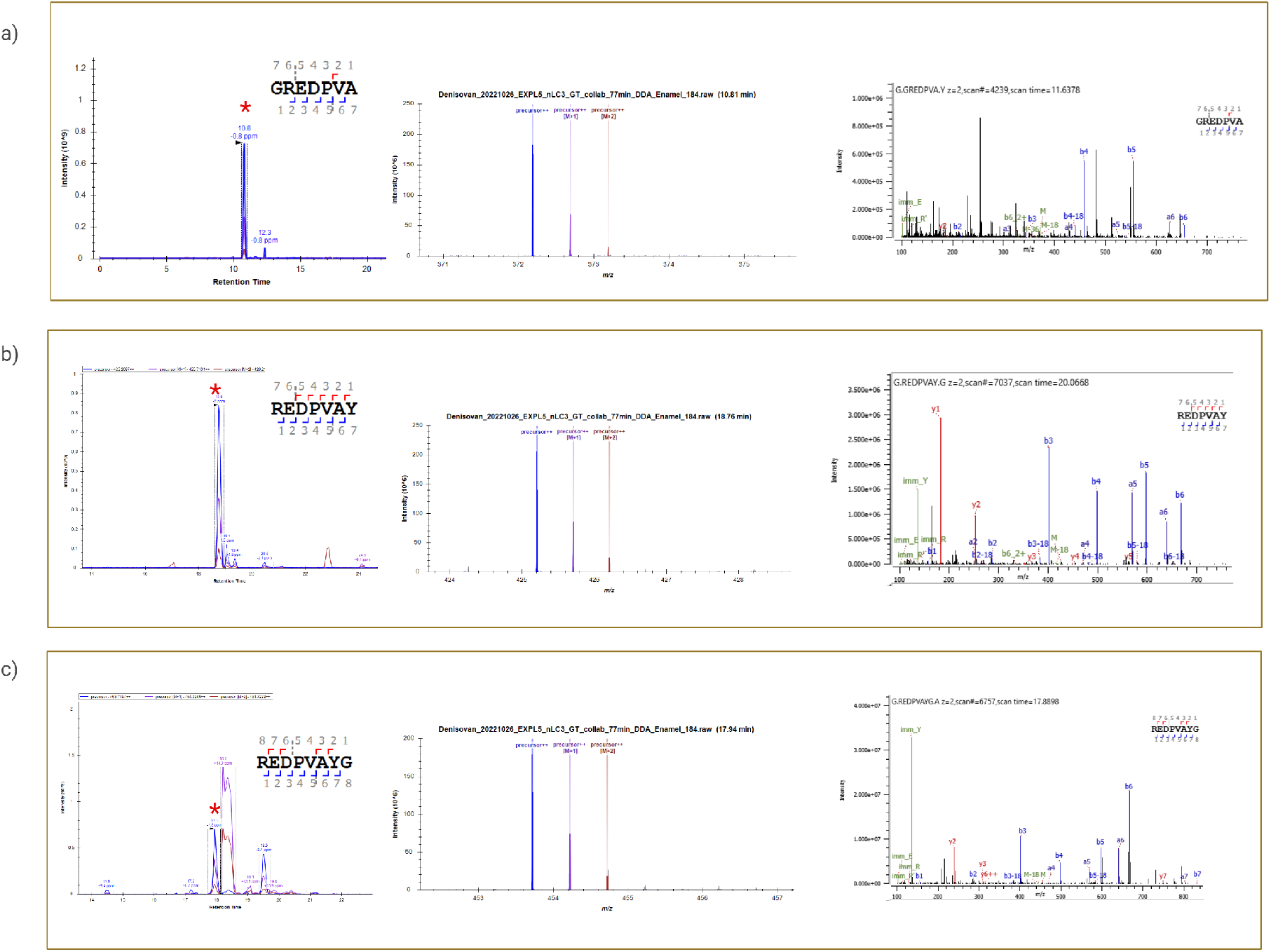
Extracted ion chromatograms (XIC) and isotopic envelopes for Denisovan-specific AMBN V273 peptides in Denisovan_184_. XIC profiles and isotopic envelope confirmation for the three highest-intensity V273-containing AMBN peptides detected in Denisovan_184_: **(A)** GREDPVA (*m/z* 372.19, charge 2+; RT 11.2 min; normalised area 9.15 ×10^9^); **(B)** REDPVAY (*m/z* 425.21, charge 2+; RT 19.44 min; normalised area 1.10 ×10^10^); **(C)** REDPVAYG (*m/z* 453.72, charge 2+; RT 19.48 min; normalised area 8.11 × 10^9^). For each peptide, the panel shows the XIC with the integrated peak area, the observed isotopic envelope, and the MS2 Byonic identification. Retention times are consistent with Byonic identification anchors from the same dataset.

**Figure 2.**
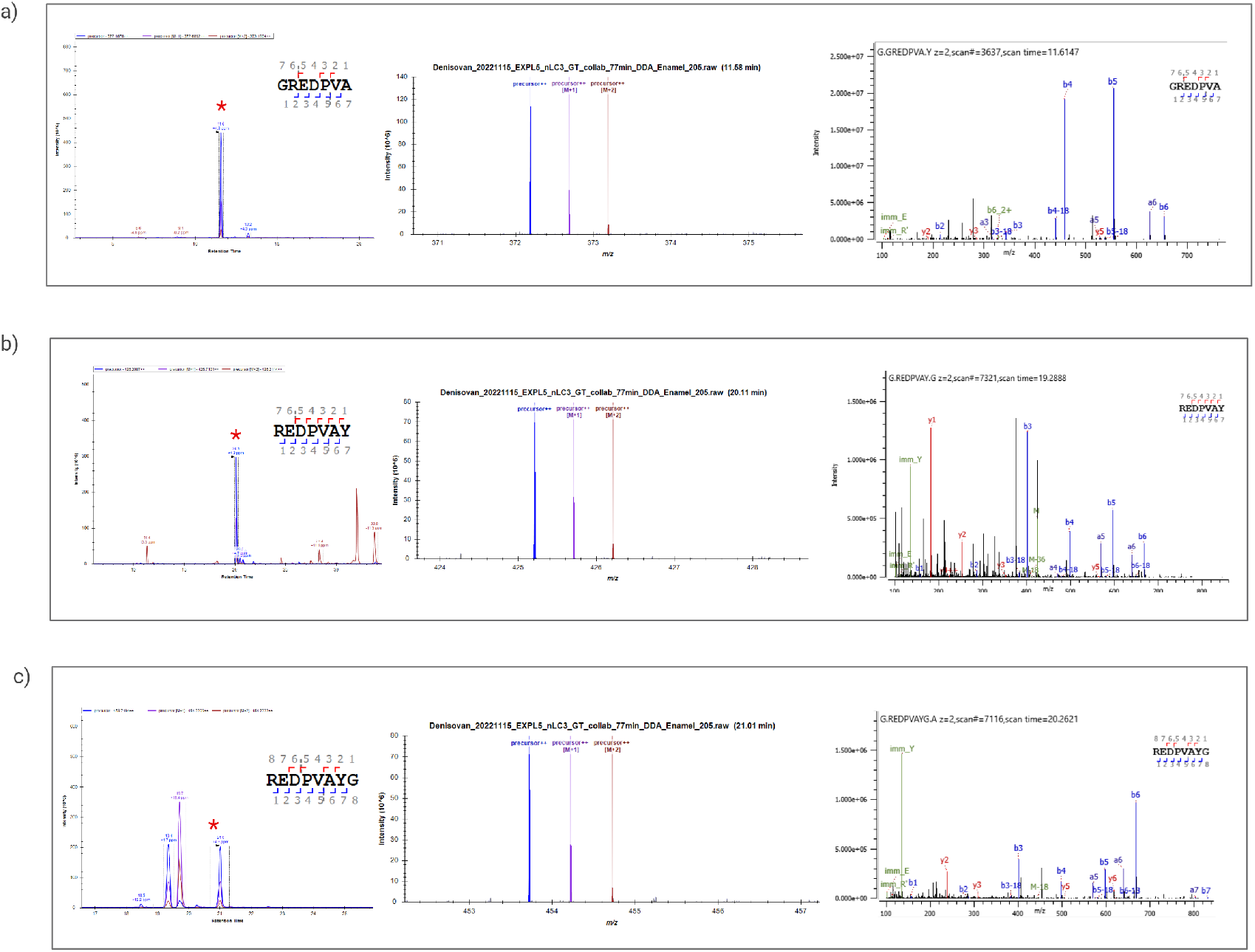
Extracted ion chromatograms (XIC) and isotopic envelopes for Denisovan-specific AMBN V273 peptides in Denisovan_205_. XIC profiles and isotopic envelope confirmation for the three highest-intensity V273-containing AMBN peptides detected in Denisovan_205_: **(A)** GREDPVA (*m/z* 372.19, charge 2+; RT 11.2 min; normalised area 3.49 × 10^9^); **(B)** REDPVAY (*m/z* 425.21, charge 2+; RT 19.44 min; normalised area 2.99 ×10^9^); **(C)** REDPVAYG (*m/z* 453.72, charge 2+; RT 19.48 min; normalised area 1.74 ×10^9^). For each peptide, the panel shows the XIC with the integrated peak area, the observed isotopic envelope, and the MS2 Byonic identification. Retention times are consistent with Byonic identification anchors from the same dataset.

**Figure 3.**
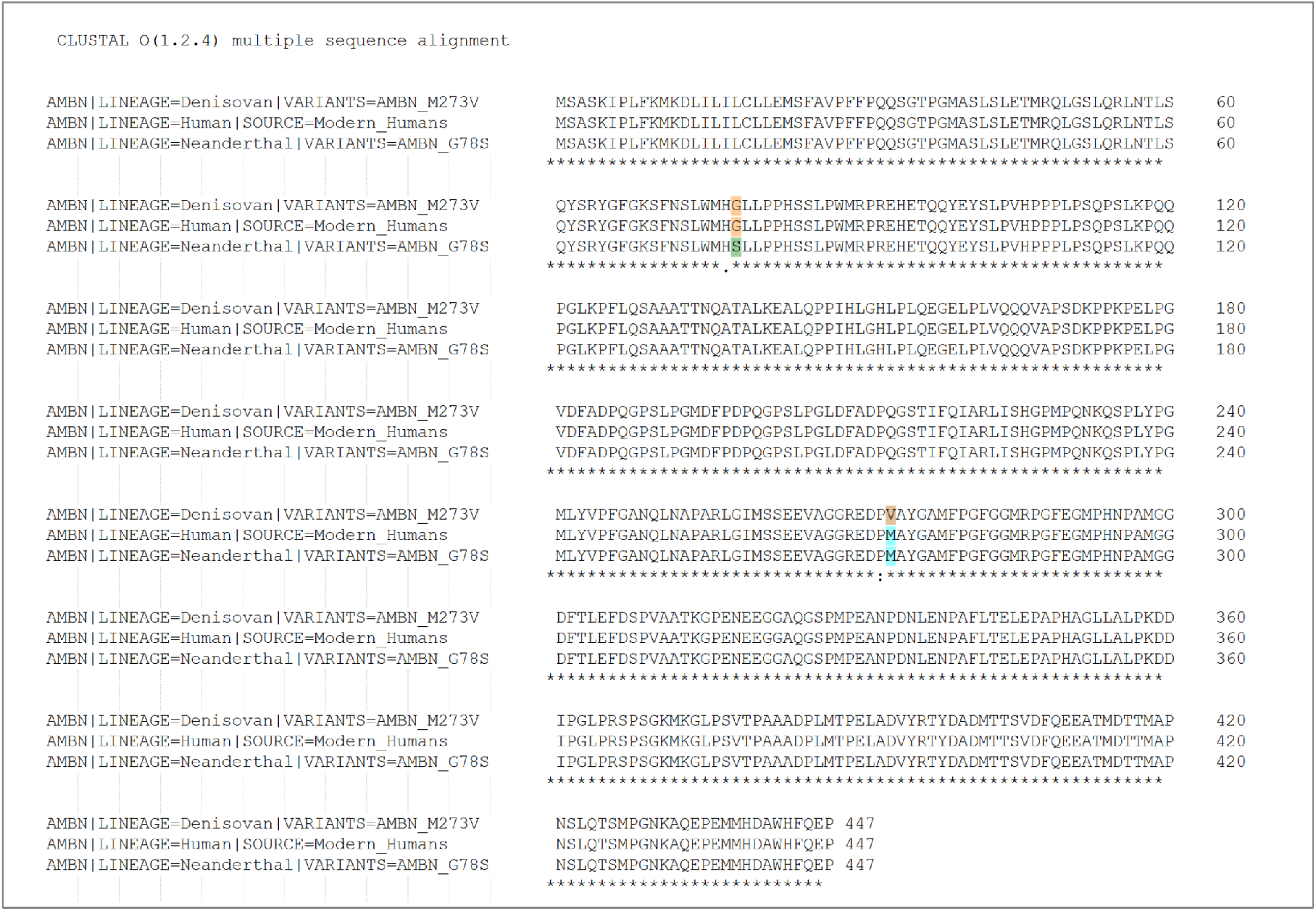
AMBN sequence alignment at positions 78 and 273 across Denisovans, Neandertals, and modern *Homo sapiens*. Partial alignment of the ameloblastin (AMBN) protein sequence centred on residues 78 and 273. Neandertal individuals (Altaï and Vindija) carry the ancestral methionine at position 273 (M273) and the ancestral glycine at position 78 (G78). The Denisovan individual from Denisova Cave carries both derived substitutions: valine at position 273 (V273; rs564905233) and serine at position 78 (S78; rs143795139). In modern *Homo sapiens*, V273 and S78 occur at extremely low allele frequencies (≈ 0.001 and ≈ 0.0001, respectively), making V273 effectively a Denisovan-diagnostic residue in the Pleistocene fossil record. Derived substitutions are highlighted in red; ancestral residues are shown in grey. Sequence data adapted from Zanollli (2017) ^10^.

**Figure 4.**
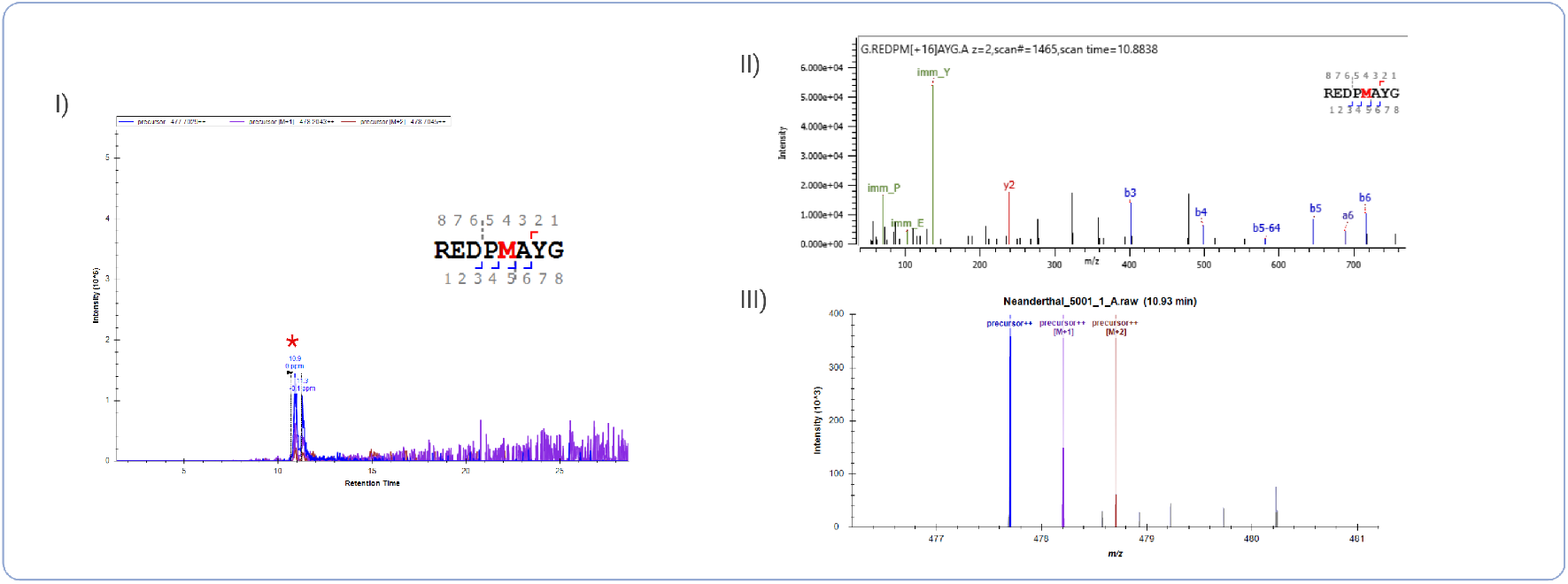
Extracted ion chromatogram (XIC) and isotopic envelope for the ancestral AMBN M273 peptide detected in the Gruta de Oliveira Neandertal . XIC profile and isotopic envelope confirmation for the sole AMBN peptide quantified in the Neandertal specimen: REDPM[+16]AYG (*m/z* 477.70, charge 2+; RT 10.9 min; normalised area 2.22 × 10^7^), carrying the ancestral methionine at position 273 with methionine oxidation (+15.99 Da). The panel shows the XIC with the integrated peak area, the observed isotopic envelope, and the MS2 Byonic identification. No V273-containing peptides (REDPVAYG and related variants) were detected in this sample, consistent with the absence of the Denisovan-specific substitution. Retention time is consistent with the Byonic identification anchor from the same dataset.

### Detection of ancestral AMBN M273 peptide in Neandertal enamel

The ancestral M273-containing peptide **REDPM[+16]AYG** (methionine oxidised; *m/z* 477.70, charge 2+; RT 10.9 min) was detected exclusively in the Neandertal specimen (see Fig. 4), yielding a normalised area of **2.22** × **107** (Table 1 and Fig. 5). This peptide did not produce a validated signal in either Penghu 1 DDA data acquisition. The mutual exclusivity of V273 signals (Denisovan only) and M273 signal (Neandertal only) across the specimens supports the taxon-discriminating specificity of the targeted assay.

**Figure 5.**
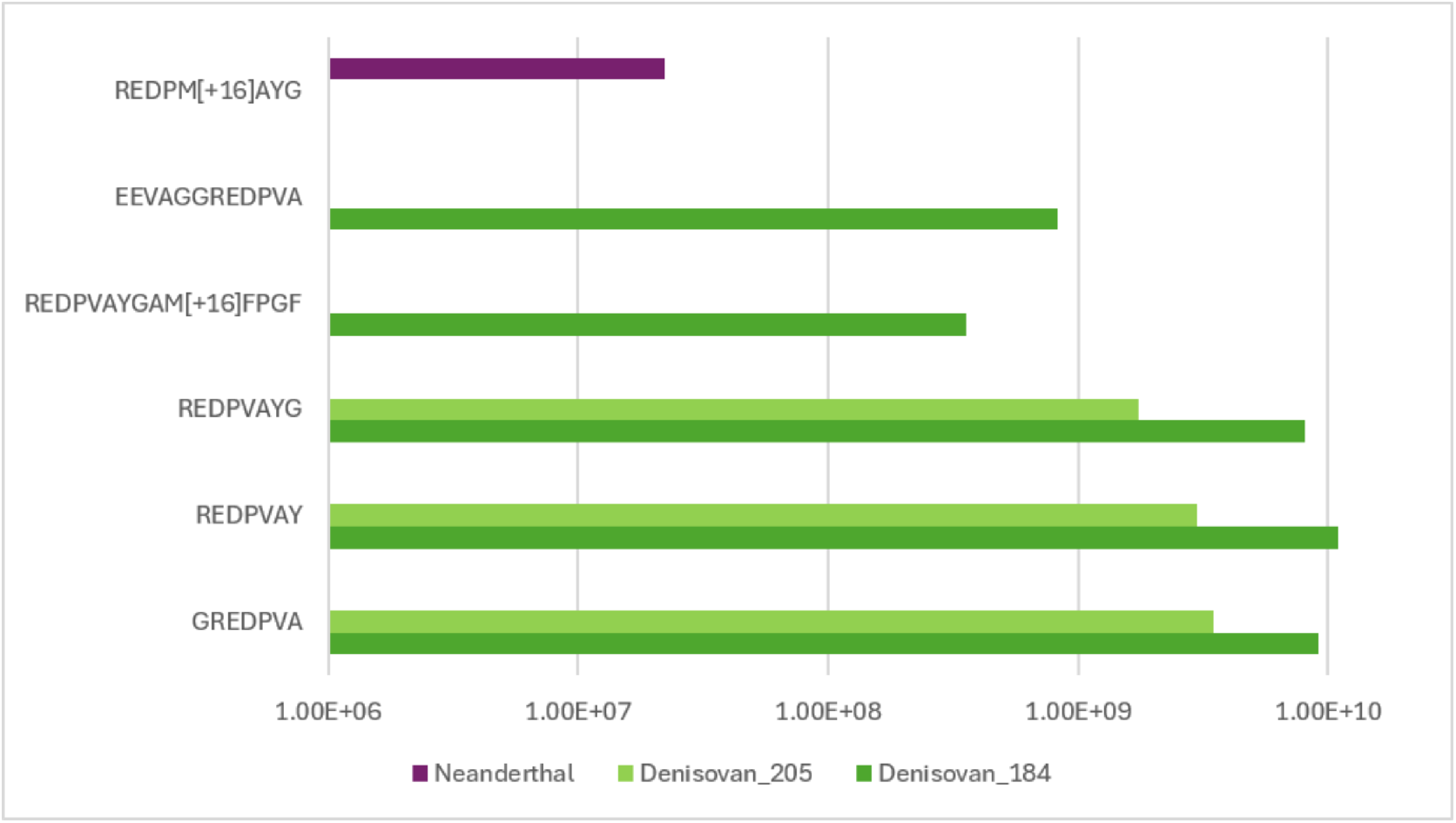
Skyline-validated AMBN peptides carrying the V273 or M273 residue across the Penghu 1 and Neandertal datasets. XIC’s are reported. ND = not detected. V273-containing peptides were detected in Penghu 1, whereas the ancestral M273-containing peptide REDPM[+16]AYG was detected only in the Gruta de Oliveira Neandertal dataset.

(*m/z* 388.71, charge 2+), were detected in both Denisovan_184_ and Denisovan_205_, confirming their identification as males, and so corroborating the results of Tsutaya et al. ^2^. SM[+16]IRPPY (1.30 × 10^8^), M[+16]IRPPY (6.46 × 10^8^), and MIRPPY (1.19 × 10^8^) were all detected in the Neandertal specimen, confirming its sex as **male**, consistent with the findings of Shaw et al. ^3^. AMELX-specific peptides (SIRPPYPSY, SIRPPY, IRPPYP) were detected in all ancient specimens, confirming enamel protein preservation across the sample set, see Fig. 6, 7 and 8 for the XIC isotopic envelop and MS/MS ID.

**Figure 6.**
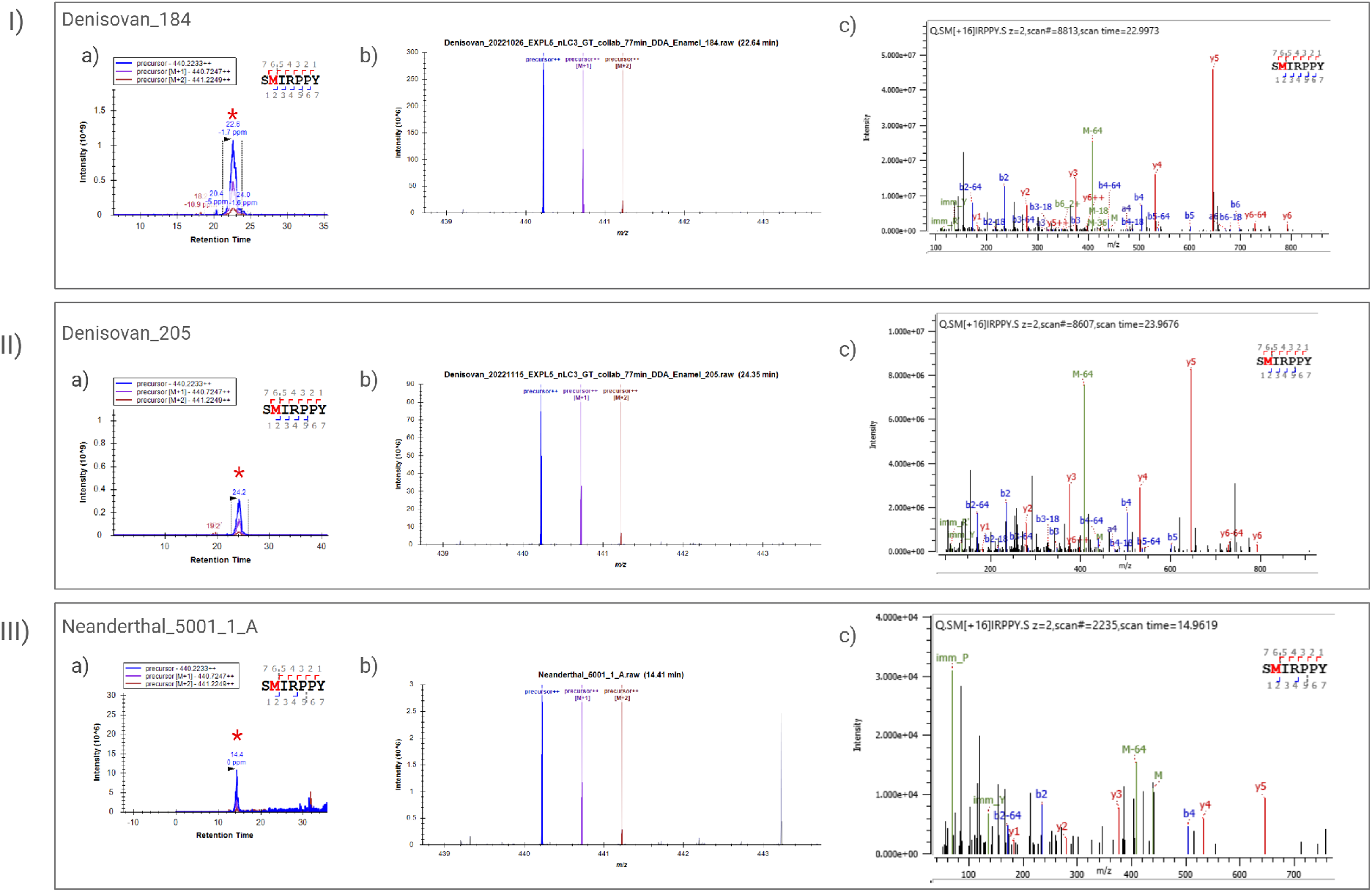
XIC profiles and isotopic envelope confirmation for the AMELY-specific peptide SM[+16]IRPPY (*m/z* 440.22, charge 2+) in the two Penghu 1 DDA data acquisitions and in the Gruta de Oliveira Neandertal dataset. Detection of this AMELY-specific peptide supports male sex assignment for both ancient individuals. **(I)** Denisovan_184_ (normalised area 7.48× 10^10^); **(II)** Denisovan_205_ (normalised area 2.49 × 10^10^); **(III)** Gruta de Oliveira Neandertal (normalised area 1.30 × 10^8^). For each panel, **(a)** shows the XIC with integrated peak area, **(b)** shows the observed isotopic envelope, and **(c)** shows the MS/MS identification by Byonic. Retention times were consistent with the corresponding Byonic identification anchors.

**Figure 7.**
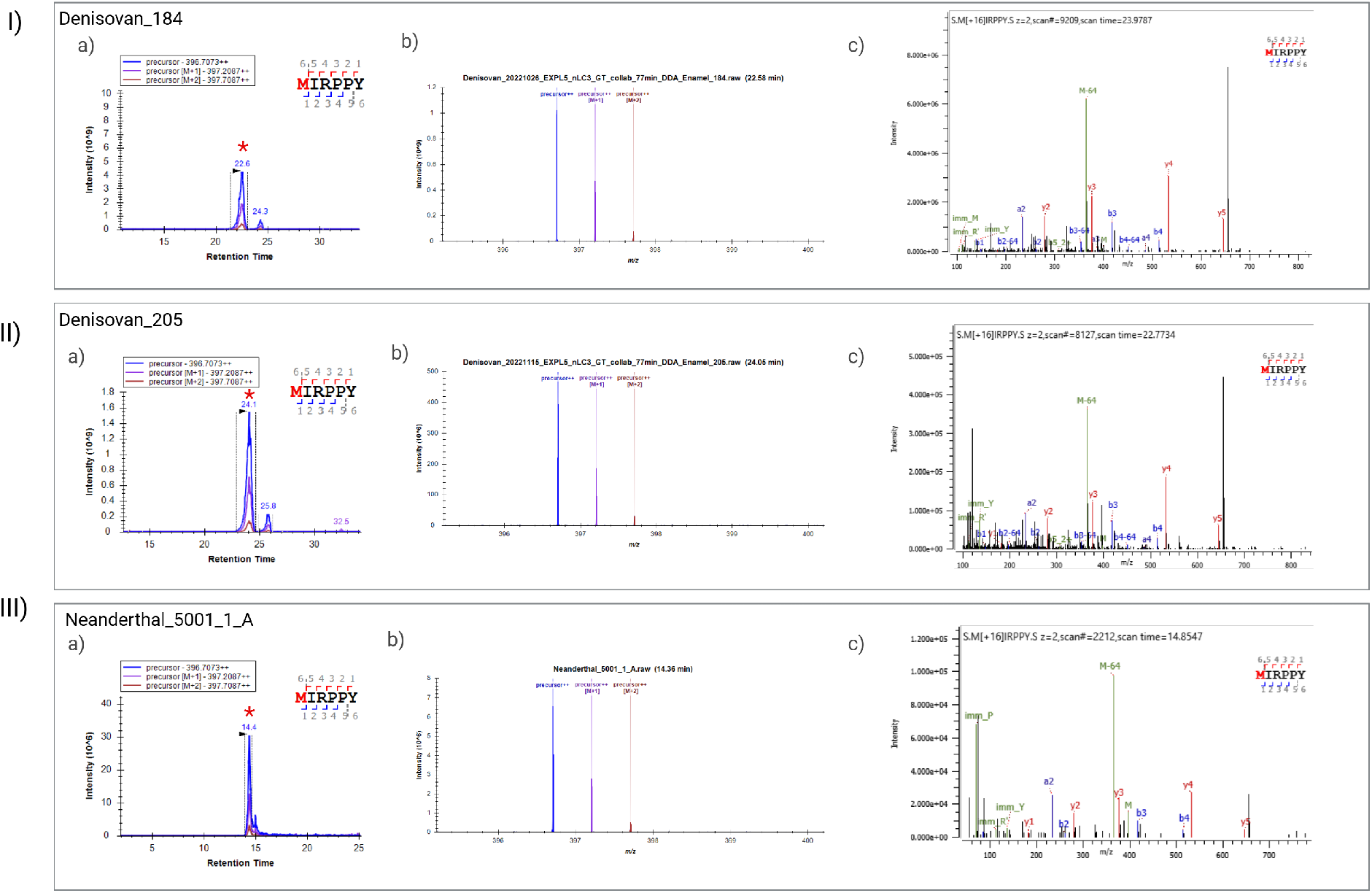
XIC profiles and isotopic envelope confirmation for the AMELY-specific peptide M[+16]IRPPY (*m/z* 396.71, charge 2+; RT 21.40 min) in the two Penghu 1 DDA data acquisitions and in the Gruta de Oliveira Neandertal dataset. Detection of this AMELY-specific peptide provides additional peptide-level support for male sex assignment in both ancient individuals. **(I)** Denisovan_184_ (normalised area 1.83× 10^11^); **(II)** Denisovan_205_ (normalised area 6.68 10^10^); **(III)** Gruta de Oliveira Neandertal (normalised area 6.46 × 10^8^). For each panel, **(a)** shows the XIC with integrated peak area, **(b)** shows the observed isotopic envelope, and **(c)** shows the MS/MS identification by Byonic. Results are concordant with Figure 6 for SM[+16]IRPPY, supporting male sex assignment for both Penghu 1 and the Gruta de Oliveira Neandertal.

**Figure 8.**
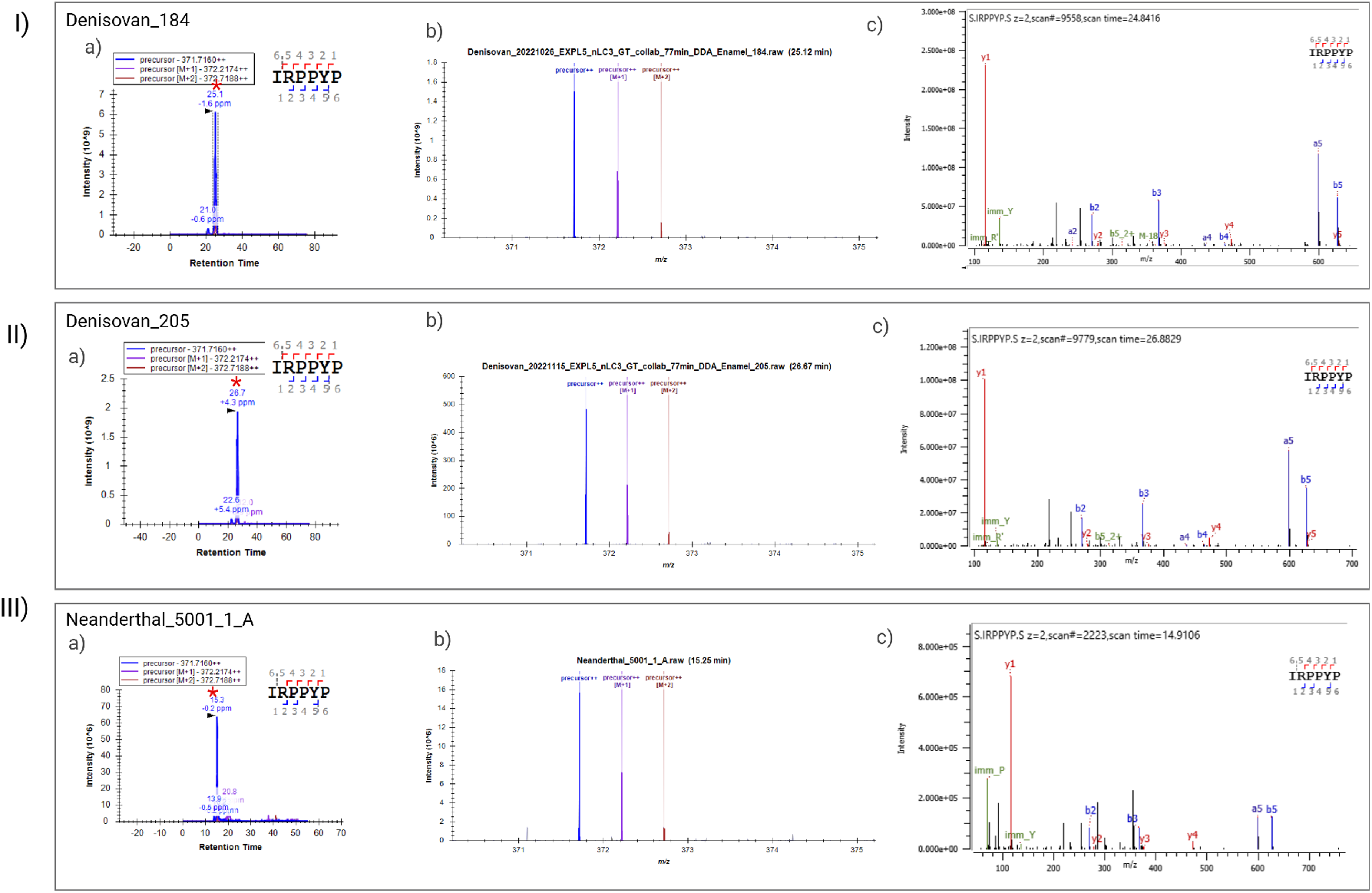
XIC profiles and isotopic envelope confirmation for the AMELX-specific peptide IRPPYP (*m/z* 371.72, charge 2+; RT 22.35 min) in the two Penghu 1 DDA data acquisitions and in the Gruta de Oliveira Neandertal dataset. Detection of this AMELX peptide confirms enamel protein preservation across the ancient datasets. **(I)** Denisovan_184_ (normalised area 3.56 × 10^11^); **(II)** Denisovan_205_ (normalised area 1.17 × 10^11^); **(III)** Gruta de Oliveira Neandertal (normalised area 1.46 × 10^9^). For each panel, **(a)** shows the XIC with integrated peak area **(b)** shows the observed isotopic envelope, and **(c)** shows the MS/MS identification by Byonic. Together with the AMELY-specific peptide evidence shown in Figures 6 and 7, these results support male sex assignment for both Penghu 1 and the Gruta de Oliveira Neandertal.

The lower absolute signal intensity for the Neandertal M273 peptide (∼360-fold below the primary Denisovan V273 signal in Denisovan_184_) is consistent with the relatively lower overall protein preservation expected for the Gruta de Oliveira specimen (∼90–92 ka BP) compared to Penghu 1, and with inherent differences in ionisation efficiency between the two peptide sequences. Qualitative presence/absence of each variant, rather than absolute area comparison across specimens, represents the primary diagnostic criterion.

### Sex determination from AMELX and AMELY peptides

AMELY-specific peptides were detected in both Penghu 1 and in the Gruta de Oliveira Neandertal dataset, supporting assay specificity for sex determination (Table 2). Specifically, three AMELY peptides, SM[+16]IRPPY (*m/z* 440.22, charge 2+), M[+16]IRPPY (*m/z* 396.71, charge 2+), and MIRPPY

**Table 2.** Amelogenin peptides used for sex determination across all specimens. Normalised peak areas reported. ND = not detected. ND^†^ = not targeted in this sample or below detection threshold. Met-ox = methionine oxidation (+15.9949 Da). Absence of AMELY signal in the modern female control confirms sex assay specificity, modern data from Kotli et. al ^13^ only for comparison.

| Peptide sequence | Protein | Denisovan<sub>184</sub><br>XIC | Denisovan<sub>205</sub><br>XIC | Neandertal<br>XIC | Modern<br>female XIC | Modern<br>male XIC |
| --- | --- | --- | --- | --- | --- | --- |
| IRPPYP | AMELX | $3.56 \times 10^{11}$ | $1.17 \times 10^{11}$ | $1.46 \times 10^9$ | ND<sup>†</sup> | ND<sup>†</sup> |
| SIRPPY | AMELX | $9.48 \times 10^{10}$ | $3.55 \times 10^{10}$ | $5.44 \times 10^8$ | ND<sup>†</sup> | ND<sup>†</sup> |
| SIRPPYPSY | AMELX | $2.39 \times 10^{10}$ | $7.47 \times 10^9$ | $1.65 \times 10^8$ | $1.77 \times 10^9$ | $7.90 \times 10^8$ |
| SM[+16]IRPPY | AMELY | $7.48 \times 10^{10}$ | $2.49 \times 10^{10}$ | $1.30 \times 10^8$ | ND | $2.49 \times 10^9$ |
| M[+16]IRPPY | AMELY | $1.83 \times 10^{11}$ | $6.68 \times 10^{10}$ | $6.46 \times 10^8$ | ND | $2.61 \times 10^8$ |
| MIRPPY | AMELY | $1.27 \times 10^{10}$ | $3.78 \times 10^9$ | $1.19 \times 10^8$ | ND | ND<sup>†</sup> |
| SMIRPPY | AMELY | $1.06 \times 10^9$ | ND<sup>†</sup> | ND | ND | $7.33 \times 10^8$ |

The deamidation of Glutamine (Q) was highly consistent across Denisovan_184_ (78.5%), Denisovan_205_ (79.4%), and the Neandertal dataset (78.7%). Asparagine (N) deamidation was 66.9% in both Penghu 1 acquisitions and 92.0% in the Neandertal dataset, although the Neandertal N estimate is based on a smaller number of N-containing residues. High deamidation rates at N and Q residues are consistent with the presence of ancient protein and serves as an internal authentication criterion, distinguishing endogenous ancient peptides from modern contaminants ^6,12^.

## Discussion

### Targeted LFQ validation as a higher evidential standard

Tsutaya et al. ^2^ reported ≥17 PSMs covering AMBN position 273 in the acid-etched fraction of Penghu 1, providing initial evidence for the V273 substitution in this specimen. Our targeted label-free re-analysis provides an independent quantitative validation of these findings. The multi-level validation strategy applied here, XIC peak integrity at consistent retention times, isotopic envelope confirmation, and MS2 ion matching (see the Skyline targeted analysis subsection in Methods), supports the assignment of each reported V273-containing peptide and reduces the likelihood of database-search misassignment. XIC analysis is particularly important for ancient specimens, where low-intensity endogenous peptide signals must be distinguished from background noise, co-eluting ions, and potential contaminant-derived features. The reproducibility of V273 detection across two independent DDA acquisitions from the same specimen, with concordant retention times and broadly consistent signal ratios for three co-detected peptides (GREDPVA, REDPVAY, REDPVAYG), substantially strengthens confidence in the endogenous ancient origin of these signals. In palaeoproteomics, replicate detection within a specimen is a key criterion for distinguishing genuine ancient peptides from contaminants or artefacts ^6^. The detection of the extended peptide REDPVAYGAM[+16]FPGF, which spans both the diagnostic position 273 and the adjacent sequence context, further corroborates the AMBN V273 assignment through a longer, more specific peptide match.

### Cross-specimen discrimination demonstrates assay specificity

A central and novel contribution of this study is the simultaneous detection of V273 (Denisovan) and M273 (ancestral) variants in two different ancient hominin specimens within a single validated Skyline workflow. The strict mutual exclusivity of these signals, five V273 peptides absent from the Neandertal; M273 present in the Neandertal and absent from both Denisovan runs, provides direct empirical evidence for the taxon-discriminating specificity of AMBN position 273 at the proteomics level (Figure 5). This cross-specimen validation was not possible from the Tsutaya et al. ^2^ study alone, which focused exclusively on the Penghu 1 specimen.

The detection of REDPM[+16]AYG in the Neandertal specimen, the methionine-oxidised form of the ancestral M273 peptide, is consistent with well-characterised post-mortem methionine oxidation chemistry in ancient proteins ^6^, and fully consistent with the taxon-specific M273/V273 distribution established by Zanolli et al. ^10^ and Froment et al. ^11^ (see Introduction). Notably, REDPM[+16]AYG was absent from both Denisovan runs, providing a true negative result that validates the V273/M273 mutual exclusivity at the targeted proteomics level.

The lower absolute signal intensity for the Neandertal M273 peptide (∼360-fold below the primary Denisovan V273 signal in Denisovan_184_) is consistent with the relatively lower overall protein preservation expected for the Gruta de Oliveira specimen (∼90–92 ka BP) compared to Penghu 1, and with inherent differences in ionisation efficiency between the two peptide sequences. Qualitative presence/absence of each variant, rather than absolute area comparison across specimens, represents the primary diagnostic criterion.

### Implications for Denisovan identification and population history

Denisovans are morphologically poorly constrained and may have occupied a broad geographic range across eastern Eurasia. Fossil specimens attributable to Denisovans have been confirmed from Denisova Cave, Siberia ^8^, the Tibetan Plateau (Xiahe mandible) ^7^, the Annamite Chain of northern Laos (Cobra Cave molar) ^9^, and now Taiwan (Penghu 1 mandible) ^2^. This geographic distribution, spanning high altitude Central Asia to low-latitude Southeast Asia, underscores the need for robust molecular identification tools applicable to specimens recovered from diverse depositional contexts where aDNA is rarely preserved.

The confirmation of AMBN V273 as a Denisovan-diagnostic proteomic marker in Penghu 1, and the characterisation of a validated targeted assay for its detection, opens practical pathways for applying this approach to other Pleistocene mandibular and dental specimens from East and Southeast Asia where Denisovan presence is suspected but aDNA is unavailable. The V273 variant is present in Denisova 3 and reaches a notably elevated frequency of ∼21% in modern Philippine populations (compared to *<*1% globally) ^2^, independently validating the Denisovan origin of this variant and reflecting documented Denisovan introgression events in island Southeast Asia ^17^. This population-level signal implies that the targeted AMBN assay could encounter V273 in modern human enamel at sites in the Philippines and nearby regions; integration with archaeological context, direct radiocarbon dating, and additional independent SAAV markers are therefore recommended for unambiguous taxonomic assignment in these geographic contexts.

Beyond M273V, Zanolli et al. ^10^ predicted AMBN G78S (rs143795139) as a Neandertal -specific variant in the Vindija 33.15 specimen. The present LFQ workflow could be extended to target G78S-containing peptides (AMBN position 78: Glycine in Denisovans and modern humans; Serine in Neandertals), enabling a multi-variant AMBN array, AMBN M273V for Denisovans, AMBN G78S for Neandertals, for distinguishing archaic hominins and modern humans in a single targeted analysis. Such a compilation would represent a significant advance in SAAV based hominin identification capability.

### Establishing a reusable targeted proteomics framework

A practical outcome of this study is the demonstration that AMBN SAAV-based hominin identification can be implemented as a standardised, reusable LFQ method with a defined targeted peptide list, precursor ion parameters, and validated retention time. This workflow is applicable to any Pleistocene enamel specimen for which DDA data are available, either from new analyses or through re-analysis of publicly archived datasets in ProteomeXchange ^1^. More broadly, this study highlights the scientific value of public proteomic repositories, where published raw data can be re-analysed with targeted approaches to test new biological questions, validate candidate markers, and extend the interpretive value of previously generated datasets.

## Conclusions

We present a targeted label-free re-analysis of published DDA data supporting the AMBN M273V substitution in Penghu 1 Pleistocene dental enamel. Five V273-containing AMBN peptides were validated in the Penghu 1 dataset by XIC, isotopic envelope confirmation, and MS2 evidence, three of which were independently detected across both DDA acquisitions. The ancestral M273 peptide REDPM[+16]AYG was detected exclusively in the Gruta de Oliveira Neandertal dataset, supporting AMBN V273/M273 discrimination between the Denisovan attributed Penghu 1 specimen and the Neandertal comparator. AMELY-specific peptides additionally support male sex assignment for both ancient individuals. Together, these results support AMBN M273V as a candidate Denisovan-diagnostic proteomic marker and establish a reproducible targeted LFQ framework for SAAV based taxonomic and nd amelogenin-based sex assessment of Pleistocene hominin enamel.

## Data Availability

Raw DDA data are available from the ProteomeXchange Consortium under accessions **PXD054412** 2 and **PXD038154** 3. The Skyline document, targeted peptide list, and annotated MS2 spectra generated in this study will be deposited in a publicly accessible repository upon publication.

## Supporting Information

Additional peptide identification results, extracted-ion chromatograms, isotopic-envelope assessments, annotated MS/MS spectra, custom protein sequence database entries, and Skyline targeted-analysis parameters (PDF and XLSX).

## Acknowledgements

I acknowledge funding from the Israel Science Foundation (grant 2075/22 to V.S.). We thank Dr. Viviane Slon and Prof. Hyla May for their support, guidance. I am also grateful to Dr. Liora Kolska Horwitz, Prof. Patricia Smith and Prof. Elisabetta Boaretto for their continued support, insightful discussions, and valuable ideas.

## Author Contributions

**P.K**.: Conceptualization, Byonic database construction, Skyline targeted analysis, figure preparation, manuscript writing.

## Competing Interests

The authors declare no competing interests.

## Notes

### Competing Interest Statement

The authors have declared no competing interest.

